# Variability of core microbiota in newly diagnosed treatment-naïve paediatric Inflammatory Bowel Disease patients

**DOI:** 10.1101/317263

**Authors:** T.G.J. de Meij, E.F.J. de Groot, C.F.W. Peeters, N.K.H de Boer, C.M.F. Kneepkens, A. Eck, M.A. Benninga, P.H.M. Savelkoul, A.A. van Bodegraven, A.E Budding

## Abstract

**BACKGROUND & AIMS:** Intestinal microbiota is considered to play a crucial role in the aetiology of inflammatory bowel disease (IBD). We aimed to describe faecal microbiota composition and dynamics in a large cohort of children with de novo (naïve) IBD, in comparison to healthy paediatric controls (HC).

**METHODS:** In this prospective study, performed at two tertiary centres, faecal samples from newly diagnosed, treatment-naïve paediatric IBD patients were collected prior to bowel cleansing for colonoscopy (t0) and 1, 3 and 6 weeks and 3 months after initiation of therapy. The microbial profiles of Crohn’s disease (CD) and Ulcerative colitis (UC) patients were compared with HC and linked to therapeutic response. Microbiota composition was analysed by IS-pro technology.

**RESULTS:** Microbial profiles of 104 new IBD-patients (63 CD, 41 UC, median age 14.0 years) were compared to 61 HC (median 7.8 years). IBD was mainly characterised by decreased abundance of *Alistipes finegoldi* and *Alistipes putredinis*, which characterize a healthy state microbial core. The classifier including these core species as predictors achieved an AUC of the ROC curve of .87. Core bacteria tended to regain abundance during treatment, but did not reach healthy levels.

**CONCLUSION:** Faecal microbiota profiles of children with de novo CD and UC can be discriminated from HC with high accuracy, mainly driven by a decreased abundance of species shaping the microbial core in the healthy state. Paediatric IBD can therefore be characterized by decreased abundance of certain bacterial species reflecting the healthy state rather than by the introduction of pathogens.

## Introduction

Crohn’s disease (CD) and Ulcerative colitis (UC) are the two main phenotypes of inflammatory bowel disease (IBD), which typically develop in the second or third decade of life, with up to 25% of patients presenting before 18 years of age or in young adulthood. Over the past era, incidence of IBD, especially CD, has increased globally, while the age of presentation has shown a downward trend (1,2,3,4). The aetiology of IBD is incompletely elucidated. Current data indicate a complex interplay between host (genetic) and environmental factors. The intestinal microbiome shape an important representative of the latter, by its capacity to affect barrier integrity and to induce an aberrant mucosal immune regulation (5,6,7,8). Mucosal immunological response in IBD is presumably not provoked by a single pathogen, but rather by intestinal dysbiosis, defined as an imbalance in the composition and function of intestinal microbes (9). Recognition of the possible role of the microbiome in the pathogenesis of IBD has spurred research addressing personalised, microbiota-based strategies to diagnose, monitor and even treat IBD by targeted manipulation of the microbial composition (10,11,12). First, however, IBD-specific microbial signatures should be identified and dynamics of the microbiota in IBD better understood (13). Presently it is unknown whether this microbial disturbance precedes or follows the inflammatory cascade leading to the typical IBD phenotype, and also whether (paediatric) IBD is primarily characterised by introduction of pathogenic bacteria or by the loss of health-related microbes (14,15,16). Microbiota-based studies in IBD based on a longitudinal, instead of a cross-sectional design, allow for more direct and integral testing of causality of microbe-host interactions and for evaluation whether microbiota composition can be used to predict and monitor therapeutic efficacy (17). Paediatric CD is of particular interest in this sense; in this population exclusive enteral nutrition (EEN) is the primary treatment for remission induction, instead of drug-mediated modification of the mucosal immunity as in adults, which is associated with major gut microbiota alterations (18,19). Detailed knowledge of microbiota changes after EEN might lead to improvement of composition and application of this therapy, for example in the prevention of relapses. Furthermore, prediction of efficacy of EEN based on microbiota composition at diagnosis could possibly prevent a subgroup of children from prescription of EEN, which carries a high burden on children. Unfortunately, data regarding temporal microbiota dynamics linked to therapeutic responses in children are scarce (13,20). Therefore, the aim of this study was to describe the microbiota composition and diversity in paediatric de novo CD and UC and their changes over time during treatment. Results are compared with those of healthy controls (HC), which were previously characterised by a core ‘signature’ microbiome (21). Additionally, we assessed whether the efficacy of EEN treatment in CD could be predicted based on microbiota composition at diagnosis.

## Materials and methods

### Subjects and design

Children younger than 18 years of age and presenting with treatment-naïve CD or UC, diagnosed according to the revised Porto criteria for paediatric IBD (22) were eligible to participate in this study. Between March 2011 and November 2014, eligible patients were enrolled in two tertiary hospitals (VU University Medical Centre and Emma children’s hospital, Academic Medical Centre) in Amsterdam, the Netherlands. Localisation and disease behaviour of IBD were classified by means of the Paris classification (23). Exclusion criteria were a diagnosis of IBD unclassified, the use of antibiotics, steroids or immunosuppressive therapy within the last month prior to inclusion, and proven infectious colitis within the last three months prior to inclusion. Patients undergoing diagnostic ileocolonoscopy and upper endoscopy for suspicion of IBD were requested to collect a faecal sample in a sterile container and store it in the freezing compartment of the refrigerator at home, prior to the bowel preparation. Disease activity at the time of diagnosis (t0), was assessed by physician’s global assessment (PGA) using a four-point scale: severe, moderate, mild and quiescent (24). Disease activity at t0 was substantiated by the assessment of C-reactive protein (CRP) and faecal calprotectin (FCP) levels. CD patients underwent a MRI enteroclysis to assess the presence and extent of small bowel involvement. All included patients were requested to collect additional faecal samples at 1 (t1), 3 (t3), 6 (t6) weeks and 3 (t12) months after initiation of the induction therapy. At t6, assessment of PGA, FCP and CRP levels was repeated in all patients. Patients were considered to be in clinical remission when PGA was scored as quiescent. All children diagnosed with CD were primarily offered 6 weeks EEN (polymeric Nutrison Standard®, Nutricia) for the induction of remission, according to the international guidelines (25). These children were instructed to collect the t6 follow-up sample just prior to reintroduction of the normal diet. In case of reluctance or lack of clinical response to EEN, oral corticosteroids were prescribed. Dose was based on patients body weight, with a maximum of daily 40 mg, followed by a tapering strategy according to standard care guidelines. As maintenance immunosuppressive therapy for CD patients, thiopurines were prescribed, which were started within the first weeks of EEN treatment. Patients with UC were prescribed aminosalicylates for both remission induction and maintenance therapy; dependent on disease severity at baseline and clinical response to aminosalicylates, corticosteroids were added.

As a control group a cohort was used consisting of 61 healthy children (HC) aged 4-18 years, whose microbiota profiles have been previously assessed and extensively described (21). The study was approved by University Ethics Committees of both participating centres.

#### Sampling

Sterile plastic containers and an information letter were provided to children and their parents, comprising instructions for collecting and storing the faecal samples. Children (or their parents) were instructed to collect approximately two grams per sample and to store these immediately in the (household) freezer at home at −20 °C. After (cooled) transport to the hospital, all faecal samples were immediately stored at −20 °C, until further handling.

### DNA extraction and sample preparation

As previously described (21, 26) DNA was extracted from faecal samples with the easyMAG extraction kit according to the manufacturer’s instructions (Biomérieux, Marcy l’Etoile, France). 100-400 mg of faeces was added to an Eppendorf tube containing 200μl of nucliSENS lysis buffer. This was vortexed and incubated shaking for 5 minutes at room temperature. After centrifugation (13000 rpm; 2 min), 100 μl of the supernatant was transferred to an easyMAG isolation container filled with 2 ml of nucliSENS lysis buffer. After incubation for 10 minutes at room temperature, 70 μl of magnetic silica beads were added. The easyMAG automated DNA isolation machine was used following the “specific A” protocol, eluting DNA in 110 μl buffer. All faecal samples were analysed by IS-pro (26).

### Microbiota analysis by IS-pro

For IS-pro, DNA samples were diluted 1:10. Amplification of IS-regions was performed with the IS-pro assay (IS-diagnostics Ltd, Amsterdam, the Netherlands) according to the protocol provided by the manufacturer (26). IS-pro differentiates bacterial species by the species specific length of the 16S–23S rDNA IS region with taxonomic classification by phylum-specific fluorescently labelled PCR primers.^14^ The procedure consists of two separate multiplex PCRs, the first one containing two different fluorescently labelled primers. One PCR amplifies the phyla *Firmicutes, Actinobacteria, Fusobacteria, Verrucomicrobia* (FAFV), the other amplifies the phylum *Bacteroidetes*. A separate PCR with a third labelled primer is performed for the phylum *Proteobacteria*. Amplifications were carried out on a GeneAmp PCR system 9700 (Applied Biosystems, Foster City, CA). After PCR, 5 μl of PCR product was mixed with 20 μl formamide and 0.2 μl custom size marker (IS-diagnostics). DNA fragment analysis was performed on an ABI Prism 3130 XL Genetic Analyzer (Applied Biosystems).

## Data analysis

### Basic IS-pro data assessment

All data was subjected to quality control and processed with the IS-pro proprietary software suite (IS-diagnostics), resulting in microbial profiles. These profiles contain three levels of information. First, the peak-colour sorts species into the different main phylum groups of the gastrointestinal tract (FAFV, *Bacteroidetes, Proteobacteria*). Second, the length of the 16S-23S rDNA IS region, expressed by the number of nucleotides, is used to identify bacteria at species level. Individual peaks reflect different bacterial operational taxonomic units (OTU), and were considered as individual features for downstream analyses. Finally, the quantity of the PCR product is displayed by the peak height, visualised as abundance, expressed in relative fluorescence units (RFU). Figures were made using the Spotfire software package (Tibco, Palo Alto, Ca, USA). Data were analysed with the standard IS-pro proprietary software suite (26). A log2 transformation was performed on peak heights for improved consistency of estimated correlation coefficients and improved detection of variation in less prominent species. A clustered heat map was constructed by generating a correlation matrix based on all log2 transformed IS profile data, followed by clustering with the unweighted pair group method with arithmetic mean (UPGMA). Within-sample microbial diversity was calculated as Shannon diversity index based using R 2.15.2 software package. Diversity was calculated both for each phylum and for overall microbial composition (by pooling the phyla FAFV, *Bacteroidetes* and *Proteobacteria* together). Variability in microbiota composition was depicted as principal coordinate analysis (PCoA), based on cosine distance measures, which was also used to assess longitudinal differences in microbial stability between IBD patients and controls from t0 to t12.

### Data processing

Further analyses required additional data processing. We were interested in detecting (a) differential abundance signatures (differential abundance of bacterial species between the groups of interest), and (b) classification signatures (which bacterial species have predictive power in demarcating the groups of interest). Our analyses focused on the t0 and t6 measurements. Only bacterial species that had at least 10 non-zero observations, with at least one non-zero observation in each group, were considered. This filtering ensures a minimum of variation (i.e., non-null observations) for meaningful calculations. The dimension of the final dataset at t0 was *n* = 148 observations by *p* = 162 bacterial features, while these dimensions for the t6 dataset amounted to *n* = 120 and *p* = 125.

### Abundance Signatures

A global test (27) was employed for assessing differential abundance signatures at t0 and t6. This test was applied as a method to identify patterns in complex high dimensional data and to discriminate between groups, providing a single *p*-value for a chosen bacterial set. The chosen bacterial set was the set of bacterial species retained after pre-processing (see Section ‘Data Processing’ above). The results were corrected for age and sex. In case of a statistically significant global test we evaluated to which bacterial species this statistically significant result could be attributed (microbial translation). Since we were interested in detecting overall differential abundance patterns, high-variance bacterial species were given a greater weight. We checked whether standardisation of the bacterial features would change the conclusions of the global tests performed; this was not the case we were interested in detecting overall differential abundancy patterns, bacterial features with more variance were given a greater weight.

The abundance profiles of the retained features were also assessed between the t0 and t6 measurements for those patients that were in remission at t6. As these data are essentially paired, and as the features display non-normality, a nonparametric alternative to the paired-samples t-test was used for this comparison, i.e. the Wilcoxon signed rank test. Asymptotic *p*-values were calculated, as exact *p*-values could not be calculated. The approach to multiple testing was by controlling the false discovery rate (FDR) at .05.

### Classification Signatures

Classification was performed in order to: (a) differentiate children diagnosed with IBD from healthy children on the basis of their microbiota at t0, and (b) for those that were diagnosed with CD, for prediction of the success of EEN at t6 based on the microbiome composition at t0. For the latter, the predictor data at t0 had to be matched with the patients treated with EEN at t6, constructing a dataset with *n* = 33 observations by *p* = 64 bacterial features.

Group-regularised logistic ridge regression (GRR) with variable selection (28) was applied. This classifier uses L2-penalties for regularisation and allows for the structural use of co-data (i.e., grouping of the independent variables to categories) in order to improve predictive performance. In this case, the co-data categories consisted of the phylum information to which each bacterial feature belonged to. The optimal penalty parameters were determined based on leave-one-out cross-validation of the model likelihood. Predictive performance was assessed by receiver operating characteristic (ROC) curves and the corresponding area under the ROC curves as produced by 5- fold cross-validation. The results were corrected for age and sex variables (i.e., these variables form the unpenalised additions to the classifier). Bacterial features were scaled, since features variability may differ substantially and those with high variability may drive the results.

## Results

### Patient population

Faecal samples were collected from 104 de novo IBD-patients (63 CD, 41 UC, median age 14.0 years) at diagnosis and at one, three and six weeks following initiation of therapy, and compared to 61 healthy controls (median age 7.8 years). Clinical remission at six weeks follow-up was achieved in 44/63 (70 %) and 29/41 (71%) of CD and UC patients, respectively. An additional faecal sample at t12 was collected in 46/104 patients (31 CD, 15 UC). Patient characteristics are given in Table 1. None of the children underwent intestinal surgery during the study period.

**Table 1.**
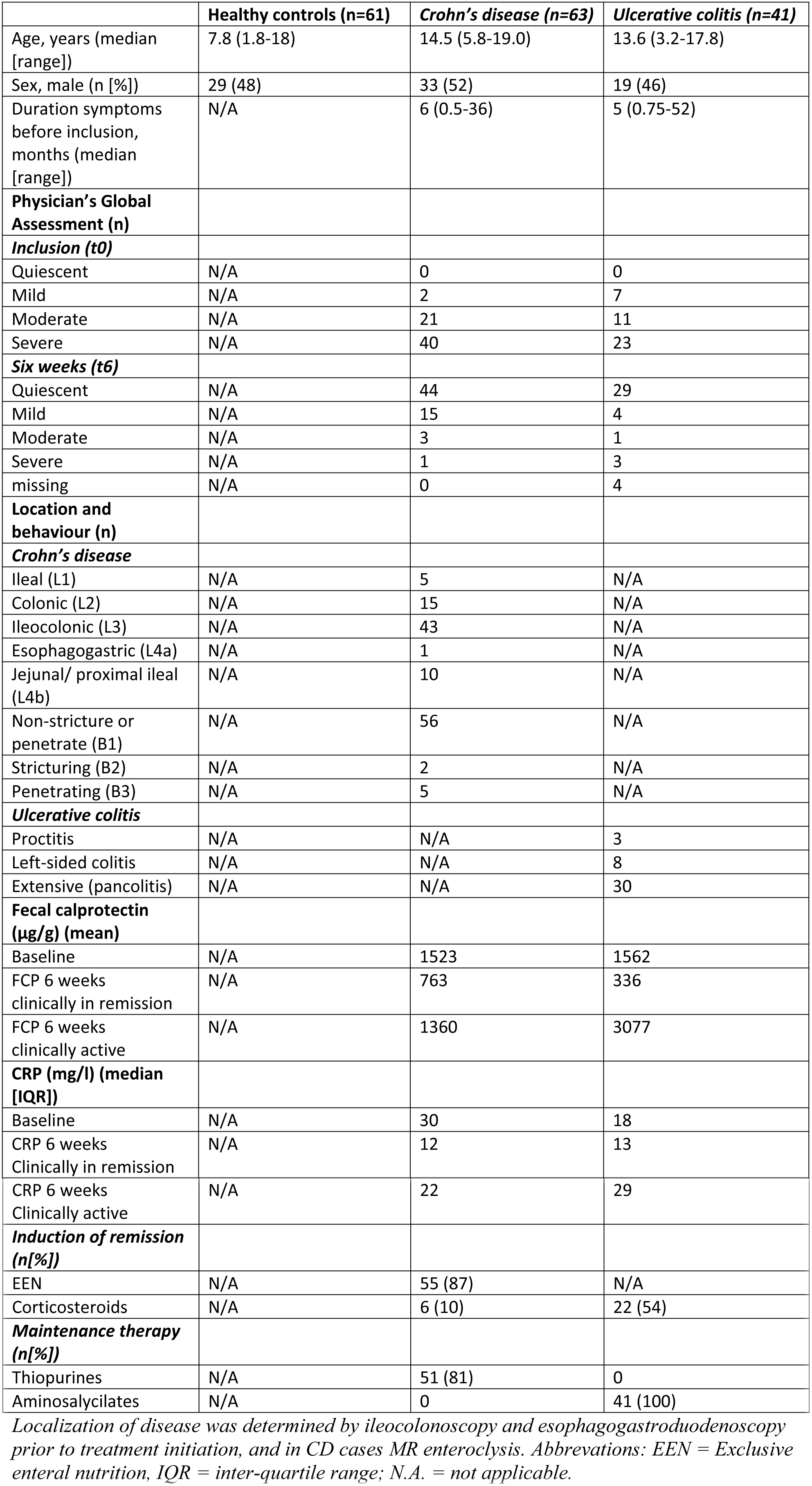
Subject characteristics of IBD subjects and controls

|  | **Healthy controls (n=61)** | **Crohn's disease (n=63)** | **Ulcerative colitis (n=41)** |
| --- | --- | --- | --- |
| Age, years (median [range]) | 7.8 (1.8-18) | 14.5 (5.8-19.0) | 13.6 (3.2-17.8) |
| Sex, male (n [%]) | 29 (48) | 33 (52) | 19 (46) |
| Duration symptoms before inclusion, months (median [range]) | N/A | 6 (0.5-36) | 5 (0.75-52) |
| **Physician's Global Assessment (n)** |  |  |  |
| **Inclusion (t0)** |  |  |  |
| Quiescent | N/A | 0 | 0 |
| Mild | N/A | 2 | 7 |
| Moderate | N/A | 21 | 11 |
| Severe | N/A | 40 | 23 |
| **Six weeks (t6)** |  |  |  |
| Quiescent | N/A | 44 | 29 |
| Mild | N/A | 15 | 4 |
| Moderate | N/A | 3 | 1 |
| Severe | N/A | 1 | 3 |
| missing | N/A | 0 | 4 |
| **Location and behaviour (n)** |  |  |  |
| **Crohn's disease** |  |  |  |
| Ileal (L1) | N/A | 5 | N/A |
| Colonic (L2) | N/A | 15 | N/A |
| Ileocolonic (L3) | N/A | 43 | N/A |
| Esophagogastric (L4a) | N/A | 1 | N/A |
| Jejunal/ proximal ileal (L4b) | N/A | 10 | N/A |
| Non-stricture or penetrate (B1) | N/A | 56 | N/A |
| Strictureing (B2) | N/A | 2 | N/A |
| Penetrating (B3) | N/A | 5 | N/A |
| **Ulcerative colitis** |  |  |  |
| Proctitis | N/A | N/A | 3 |
| Left-sided colitis | N/A | N/A | 8 |
| Extensive (pancolitis) | N/A | N/A | 30 |
| **Fecal calprotectin (µg/g) (mean)** |  |  |  |
| Baseline | N/A | 1523 | 1562 |
| FCP 6 weeks clinically in remission | N/A | 763 | 336 |
| FCP 6 weeks clinically active | N/A | 1360 | 3077 |
| **CRP (mg/l) (median [IQR])** |  |  |  |
| Baseline | N/A | 30 | 18 |
| CRP 6 weeks Clinically in remission | N/A | 12 | 13 |
| CRP 6 weeks<br>Clinically active | N/A | 22 | 29 |
| **Induction of remission**<br>**(n[%])** |  |  |  |
| EEN | N/A | 55 (87) | N/A |
| Corticosteroids | N/A | 6 (10) | 22 (54) |
| **Maintenance therapy**<br>**(n[%])** |  |  |  |
| Thiopurines | N/A | 51 (81) | 0 |
| Aminosalcilates | N/A | 0 | 41 (100) |

*Localization of disease was determined by ileocolonoscopy and esophagogastroduodenoscopy prior to treatment initiation, and in CD cases MR enteroclysis. Abbreviations: EEN = Exclusive enteral nutrition, IQR = inter-quartile range; N.A. = not applicable.*

## Microbiota analysis

### Microbial diversity

An overview of diversity indices per phylum for CU, CD and controls, at t0 and t6 is depicted in Fig 1A. Differences over time were limited, with a slight, non-significant, increase in FAFV diversity being the most outspoken observation. In Fig 1B, Shannon diversity indices per phylum over time (t0,t3,t6) are depicted for CD children on EEN. No significant differences in diversity indices were observed between the different time-points, both in clinical responders as in non-responders to EEN.

**Figure 1A.**
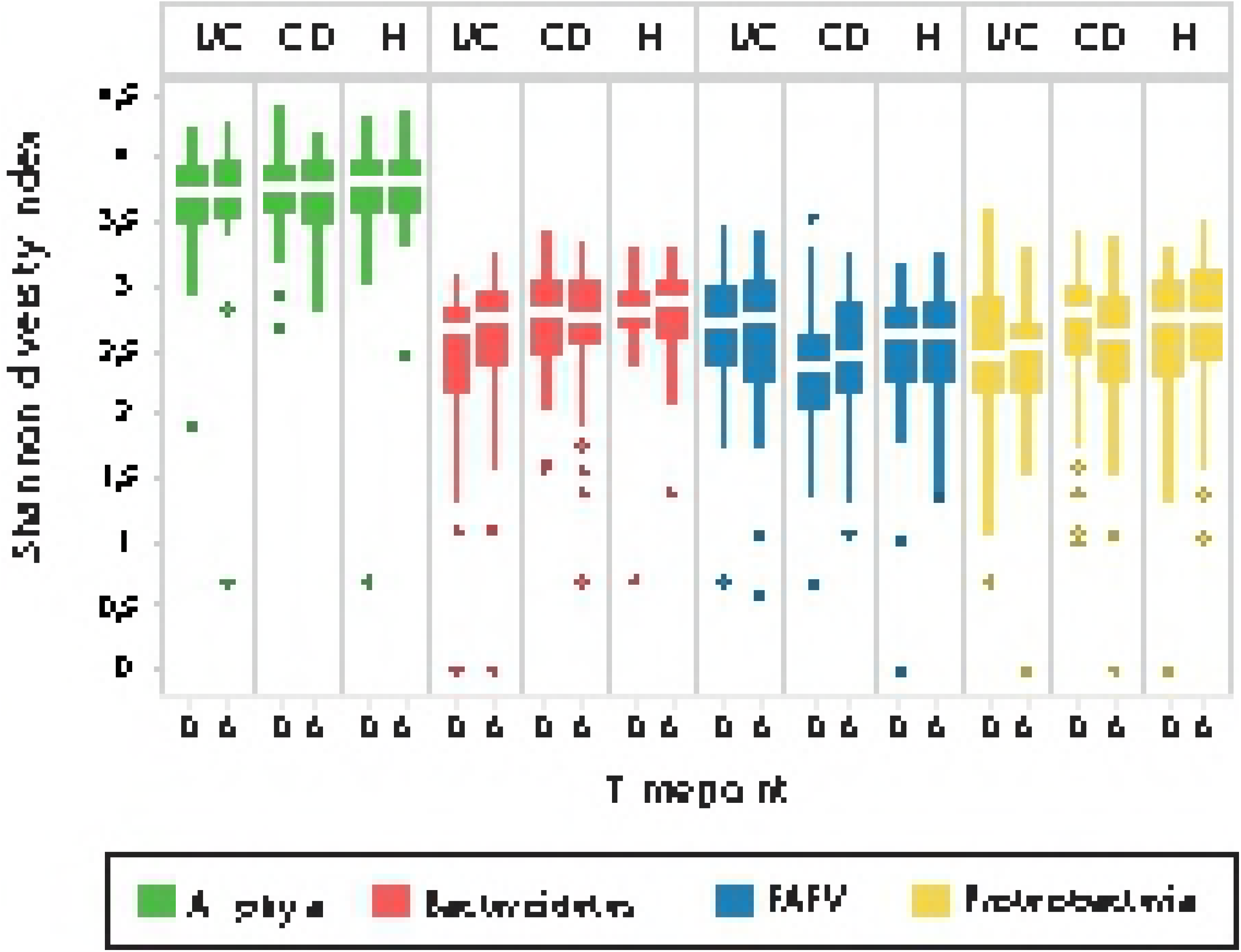
Shannon diversity indices per phylum at baseline and at six weeks follow up, during which period IBD children were treated and majority came into remission. Differences are limited, with a slight increase in *Firmicutes, Actinobacteria, Fusobacteria, Verrucomicrobia* (FAFV) diversity in CD being the most outspoken effect.

**Figure 1B.**
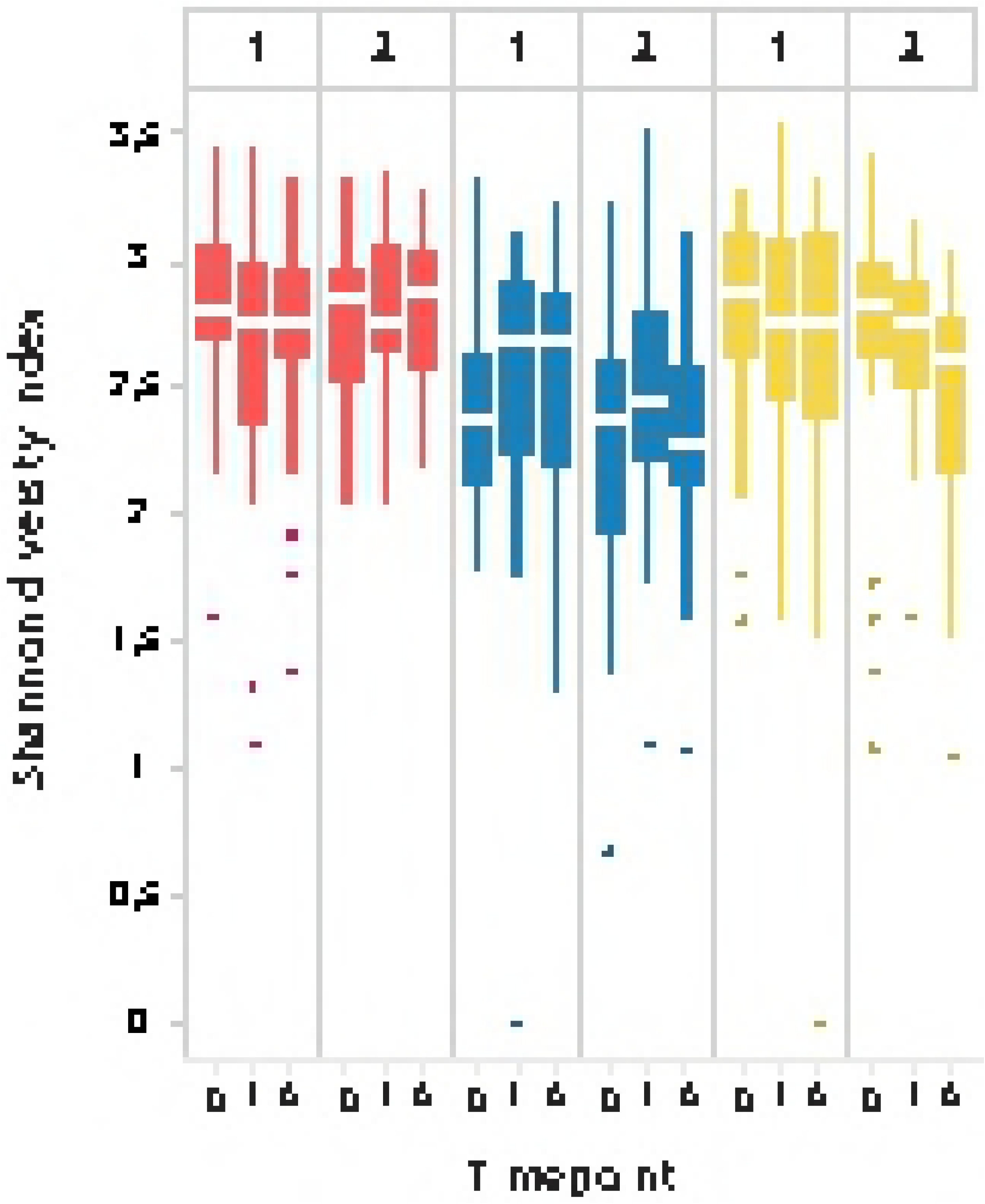
Shannon diversity indices per phylum at baseline (0) and at one and six weeks follow up for children with CD receiving exclusive enteral nutrition (EEN). No significant differences in diversity indices were observed during EEN course, both in clinical responders (1) as in non-responders (2).

### Disease-specific microbiota signatures

A clustered heatmap based on all baseline (t0) samples from all individuals showed no disease-specific clustering, indicating that, based on this unsupervised statistical approach, UC, CD and HC could not accurately be discriminated by their microbial profiles, although some disease-specific clustering was observed (Fig 2). Main characterizing feature of these subclusters was the absence of certain *Bacteroidetes*, which are abundant in healthy controls.

**Figure 2.**
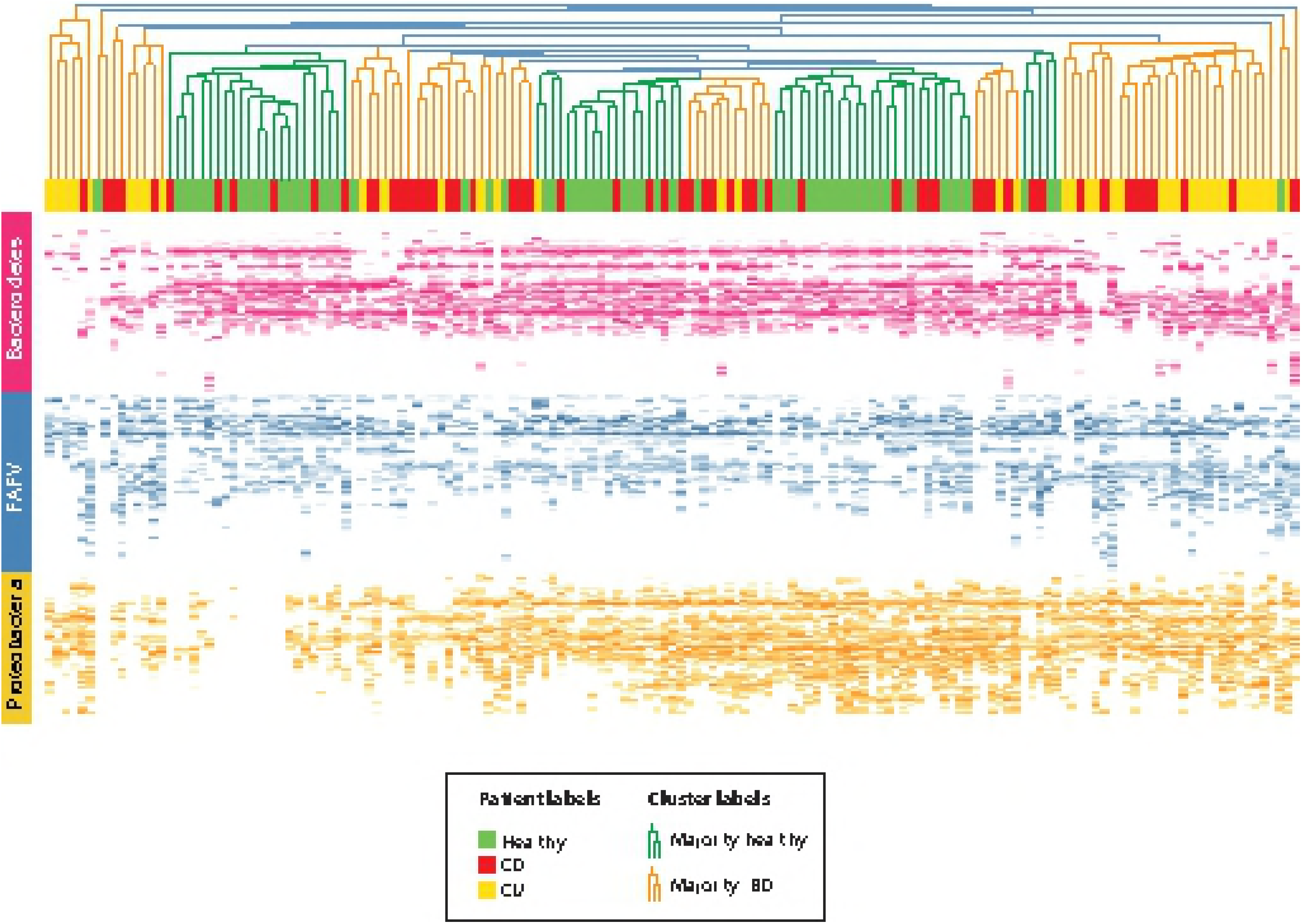
Clustered heat map of ISpro data of all baseline (t0) samples of children with ulcerative colitis (UC), Crohn’s disease (CD) and healthy controls (HC). Individual subjects are shown on the X axis; UC in yellow, CD in red, HC in green. On the Y axis, IS fragment lengths are expressed in number of nucleotides, which correspond with bacterial strain type (OTU). Blue OTUs represent *Firmicutes, Actinobacteria, Fusobacteria, Verrucomicrobia* (FAFV), red OTUs represent *Bacteroidetes* and yellow OTUs represent *Proteobacteria*. The intensity of these colors reflect the relative dominance of each indicated OTU. Some disease-specific clustering was observed: groups largely defined by the presence of children with IBD can be found at either edge of the dendrogram. Main characterizing feature of these clusters is the absence of certain *Bacteroidetes*, which are abundant in healthy controls. To some degree, further separation of IBD from healthy can be observed which are otherwise defined.

PCoA analysis of baseline samples (t0) revealed modest separation between IBD profiles and control profiles (Fig 3); this discrimination was most profound for the phylum *Bacteroidetes* (Fig 3B). However, UC and CD could not be discriminated. Intra-individual compositional stability patterns from baseline to t6, calculated based on longitudinal dissimilarity between microbial profiles, were higher in HC compared to both CD and UC (Suppl Fig 1).

**Figure 3.**
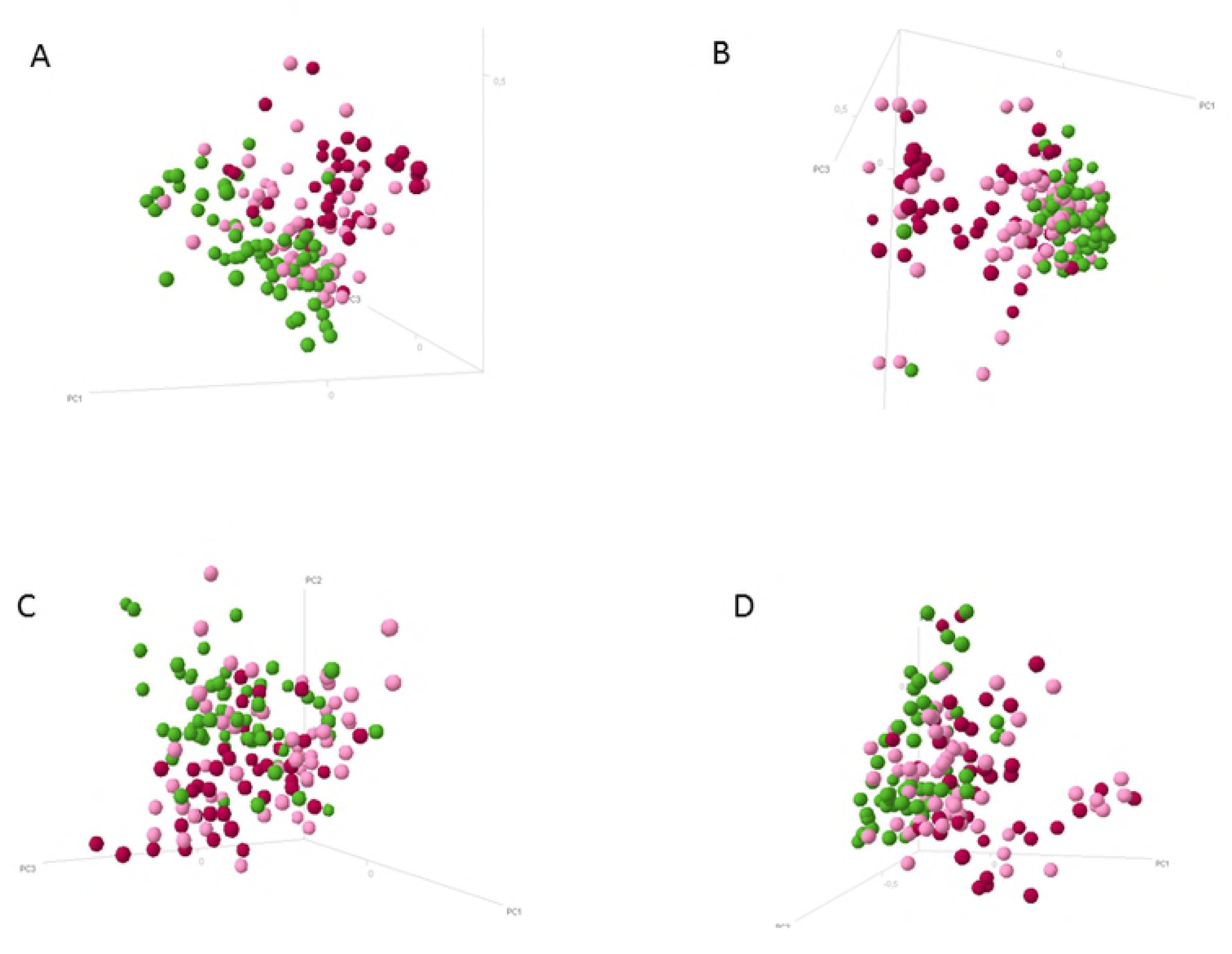
Principle coordinate analysis depicting microbial profiles of healthy controls (green), Crohn’s disease (pink) and ulcerative colitis (red) at baseline (t0), for all phyla together (3A), and per phylum (3B: *Bacteroidetes*; 3C: *Firmicutes, Actinobacteria, Fusobacteria, Verrucomicrobia* (FAFV); 3D: *Proteobacteria*). It can be seen that samples from healthy subjects can be broadly discriminated from those of IBD patients. Ulcerative colitis patients showed more separation from healthy controls than the Crohn’s disease samples. Discrimination between ulcerative colitis and controls is more profound than discrimination between Crohn’s disease and controls, in particular while taking all phyla together into account (3A). There is no apparent discrimination between ulcerative colitis and Crohn’s disease samples, for all phyla together or per phylum. Furthermore, inter-individual variation within CD subjects was higher compared to healthy controls and ulcerative colitis, in particular within the phylum *Bacteroidetes*. 3A: PCoA IBD versus controls, **all phyla** 3B: PCoA ulcerative colitis versus Crohn’s disease versus controls, **Bacteroidetes** 3C: PCoA ulcerative colitis versus Crohn’s disease versus controls, **FAFV** 3D: PCoA ulcerative colitis versus Crohn’s disease versus controls, **Proteobacteria**

### Differential abundance analysis

#### Baseline (t0)

Bacterial abundances per phylum over time, for a period of six weeks, are shown in Fig 4. The abundance of Proteobacteria declined over time for both UC and CD. Furthermore, an increase in FAFV abundance was observed for UC, concomitant to the increase in diversity.

**Figure 4.**
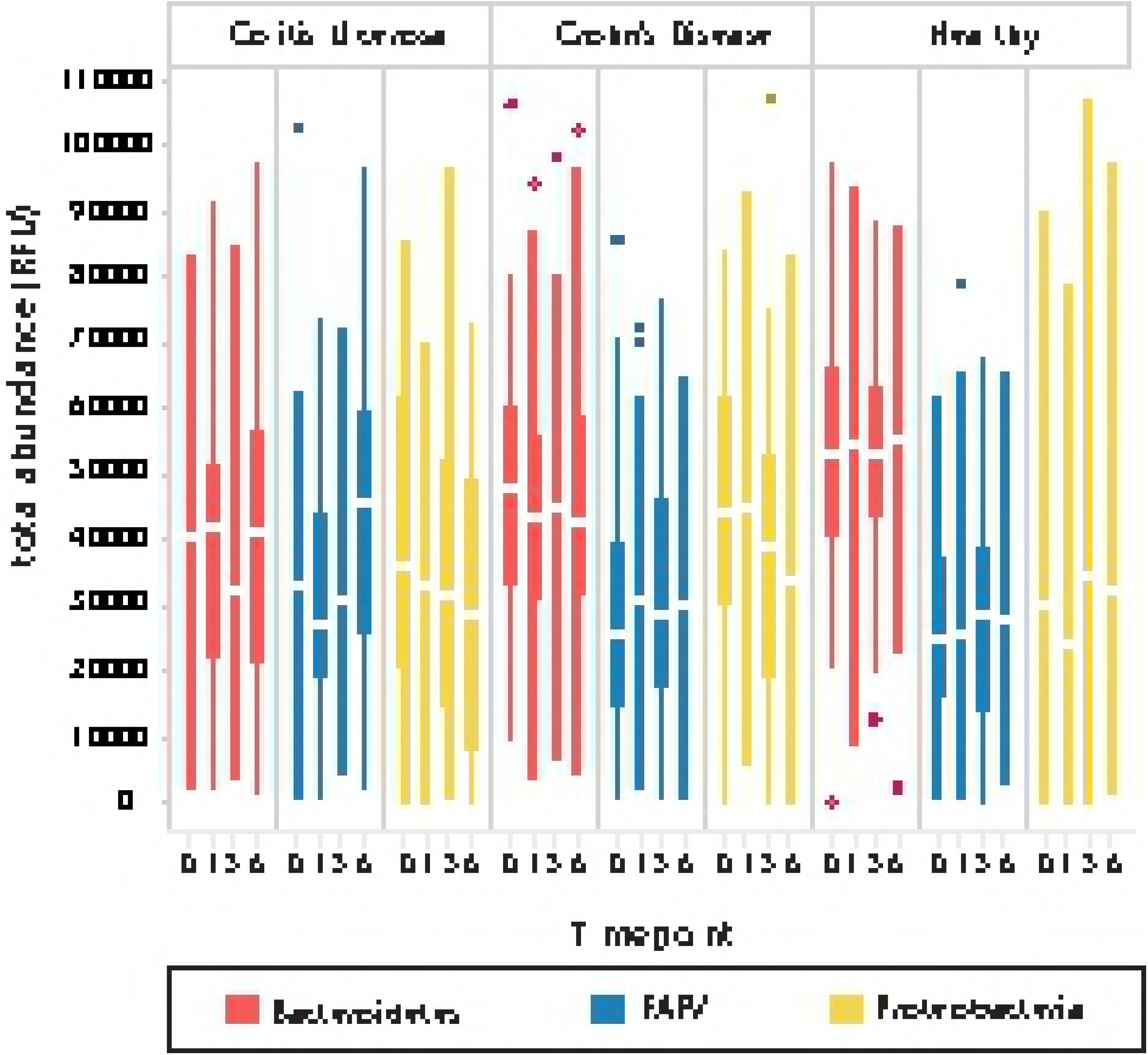
Bacterial abundances over time (baseline,1,3,6 weeks) during which period IBD children were treated. It is apparent here that the Proteobacteria abundance declines over time for both UC and CD. Also, an increase in FAFV abundance can be seen for UC, concomitant to the increase in diversity that was observed.

The global test indicated that the overall bacterial abundance profile was markedly different between healthy subjects and IBD patients (*p* < 1e-12), which could especially be attributed to the species *Alistipes putredinis* and *Alistipes finegoldi*. These species had a lower abundance in the IBD group compared to the healthy controls. Also *Escherichia coli, FAFV spp IS244, and Prevotella spp IS439* showed a significant different prevalence per disease phenotype as compared to controls at t0. Abundance of these species over a six weeks follow up period are shown in Fig 5. Phenotype-specific analysis showed that overall bacterial profiles differed between UC and CD (p = .00054); *Bacteroides fragilis* and *Akkermansia muciniphila* were decreased in UC and *Alistipes putredinis* in CD.

**Figure 5.**
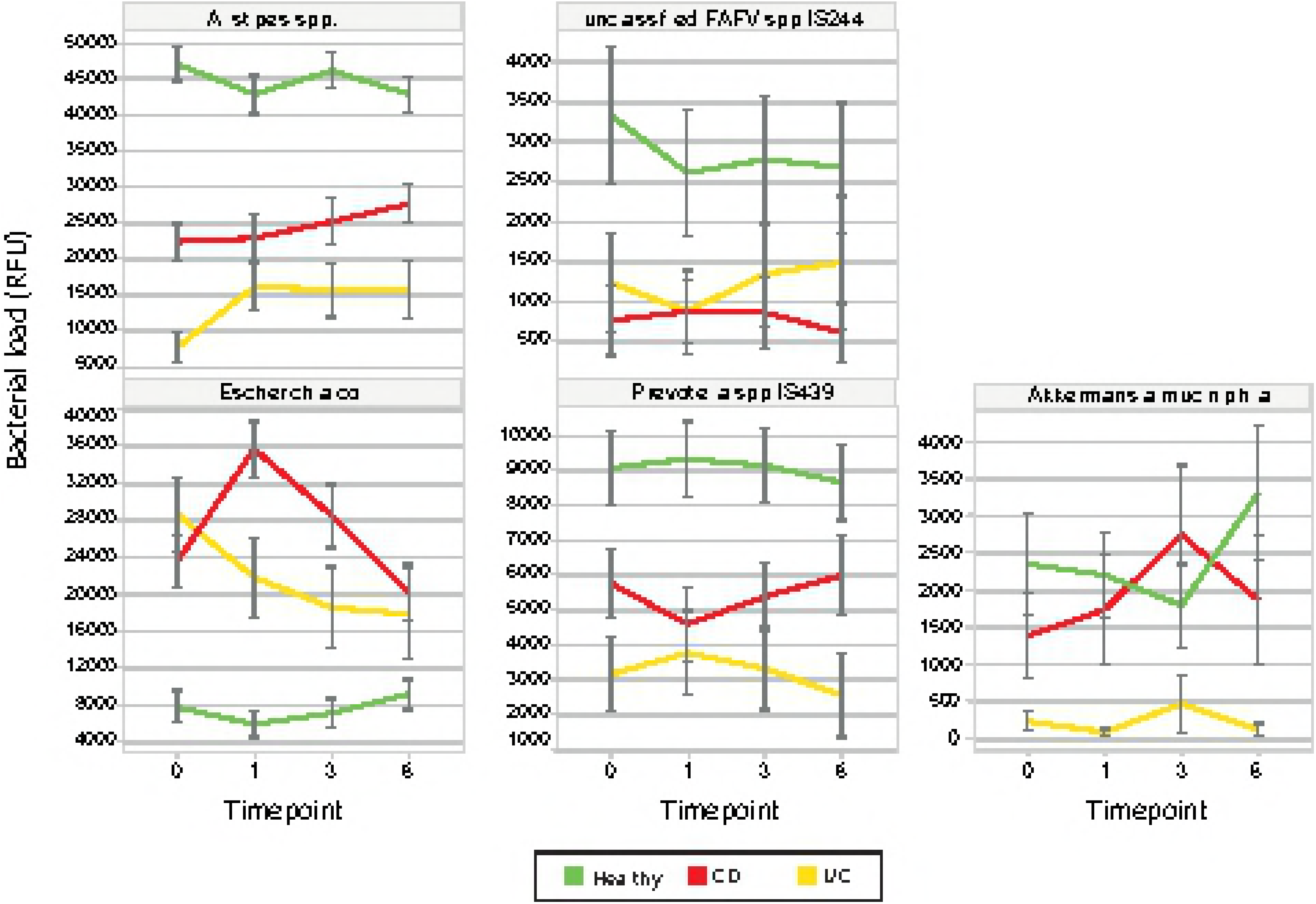
Depicted are abundances of five species with a significantly different expression per disease over time (from baseline to six weeks). The core Alistipes species (*A. finegoldii* and *A. putredinis* are taken together here, upper left) show significant lower expression in UC and CD than in HC. During treatment, levels rise towards, but do not reach, normal state. *Escherichia coli* levels (lower left) are significantly higher in UC and CD than in HC and show a marked decline (after an initial increase at week one in CD) towards normal state during treatment. Other noteworthy species are unclassified *FAFV spp IS244*, and *Prevotella spp IS439*. These species too are underrepresented in UC and CD, but do not show a concerted change during therapy. Finally, *Akkermansia muciniphila* was a special case, as it was almost absent in the entire UC group, yet showed similar abundance for CD and HC. There was no apparent effect in its presence during treatment.

### Six weeks (t6)

At six weeks follow up, UC patients who had achieved clinical remission could not be distinguished from CD patients in remission by global test. Furthermore, remission IBD microbiota profiles at t6 were not different from t6 active profiles. However, the overall abundance profiles of UC and CD (including both active and remission samples) differed markedly from the overall abundance profile in the healthy group (*p* = .00019, and *p* = .00026, respectively). The most discriminative species in these latter tests was *Eubacterium ventriosum*, which was less present in the IBD group relative to the healthy group.

### Longitudinal microbiota analysis (t0 versus t6)

Bacterial species differential abundance was assessed between t0 and t6 for patients in remission. However, none of these survived multiple testing corrections. Interestingly, the top features were those associated with the healthy state at t0. These species tended to have increased abundance in remission state. Thus, in IBD in remission, species associated with a healthy state tended to repopulate the fecal composition at t6.

### 12 weeks (t12)

Microbial course from t6 to t12 showed no clear pattern of abundance regain through time. However, sample size (n=46) was found inadequate for testing.

### Disease classification signatures

#### Baseline (t0)

A group-regularised logistic ridge regression (GRR) was used to evaluate whether IBD patients could be distinguished from controls based on their bacterial profiles. The penalty multipliers indicated that the co-data (phylum information) was informative: multipliers amounted to 0.88 for *Bacteroidetes*, 0.90 for FAFV, and 1.45 for *Proteobacteria*. This implied that *Proteobacteria* were the least informative in classification, while *Bacteroidetes* packed most discriminative power. Post-hoc variable selection resulted in 17 features, corresponding with12 different bacterial species (Table 2). All features implicated in the global test for t0 were among the top predictors in the classification signature. The AUC of the ROC curve for this model amounted to approximately .87, indicating good predictive performance for this model (Suppl. Fig 2).

**Table 2.** Post-hoc variable selection results in 17 selected features deriving from 12 bacterial species

| Phylum | Species | IS-pro peaks (nt) | Correlated with health (+) or disease (-) |
| --- | --- | --- | --- |
| Bacteroidetes | Alistipes finegoldi (core) | 230/396/400/407 | + |
| Bacteroidetes | Prevotella spp. (core) | 439 | + |
| Bacteroidetes | Alistipes putredinis (core) | 235/236 | + |
| Bacteroidetes | Bacteroides fragilis (core) | 534 | + |
| Bacteroidetes | Prevotella spp. | 577 | + |
| Bacteroidetes | Odoribacter splanchnicus (core) | 307 | + |
| Firmicutes/ FAFV | Coprothermobacter proteolyticus | 244 | + |
| Firmicutes/ FAFV | Unclassified Firmicute (core) | 541 | - |
| Firmicutes/ FAFV | Lachnospiraceae incertae sedis | 491/605 | + |
| Firmicutes/ FAFV | Streptococcus mitis group | 297 | - |
| Firmicutes/ FAFV | Faecalibacterium prausnitzii (core) | 322 | - |
| Proteobacteria | Enterobacteriaceae spp. | 889 | + |

*FAFV* = *Firmicutes*, *Actinobacteria*, *Fusobacteria*, *Verrucomicrobia*

### EEN treatment success prediction

To test the prediction of EEN treatment success in CD, as evaluated at t6, based on the microbiota composition at t0, GRR was employed. The AUC of the GRR with post-hoc variable selection amounted to .55. The AUC for the GRR without selection approximately amounted to .64. Hence, the predictive power of t0 microbial profiles with respect to EEN success at t6 was limited.

## Discussion

In this study, we examined microbiota composition and dynamics in naïve paediatric IBD. We described microbiota composition and diversity linked to disease activity in 104 de novo IBD patients compared to 61 healthy controls. Microbial profiles of IBD subjects were characterised by decreased abundance of signature species representing a microbial core in healthy state, in which *Bacteroidetes* dominate, while these core species tended to increase in abundance in remission state.

Most studies on microbiota composition in paediatric IBD have described profound differences between de novo UC, CD and controls (20,29,30,31,32). However, most reported differences are contradictory, and presumably may be explained by differences in study design, substrate collection and microbiota detection techniques. Despite apparent differences in outcome, shared observations included decreased bacterial diversity and an increase of intestinal mucosa-adhesive microbes in IBD subjects (30,31,32). Due to specific study outcomes, hypotheses regarding protective or pathogenic effects of particular species in IBD aetiology have emerged. For example, a protective role has been suggested for *Faecalibacterium prausnitzii* in paediatric CD, based on observations of its decreased abundance in affected subjects and capacities to produce proteins with anti-inflammatory properties (5,33,34). *F. prausnitzii* is therefore considered a biomarker of the healthy state. However, this presumptively protective capacity has been challenged by observations that the abundance of *F. prausnitzii* was increased in active disease and declined during EEN despite clinical improvement in CD patients (35,36). We also observed an increased abundance of *F. prausnitzii* in active disease compared to healthy state. In a cross-sectional study by Gevers and colleagues, microbial communities of 447 children with de novo CD could be differentiated with high accuracy from those of controls (AUC 0.85) (32). This microbial signature was best observed in ileal mucosa, but could not be detected in faecal samples. It was therefore suggested that faeces might be less indicative than mucosal samples to differentiate IBD patients from controls. In the present study we were able to obtain high accuracy (0.87) for correct sample classification based on microbiota from stool samples of IBD patients and controls. A possible explanation for the apparent discrepancy with the observations by Gevers et al. might be the different detection and collection methods. The use of different microbiota detection techniques may affect the outcome, preventing reliable comparison between studies (37), whereas handling, collection and storage of faecal samples are also important in the analytical outcome (38). Here, we have used IS-pro, a standardized non-selective detection method with the capacity to generate results within several hours, enforcing its potential to be applied in clinical practice, while generating comparable results to next-generation sequencing techniques (21,26). Another explanation for correct classification based on faecal samples might be statistical methodology. For example, when applying log-transformation and adapted statistical approaches on the original dataset of Gevers et al, ileal mucosal samples and faecal samples turned out to have equal predictive power (39). In the present study, acceptably sensitive classification or characterization could also not be achieved by unsupervised approaches, such as clustering and PCoA, while application of a supervised technique like group-regularised logistic regression analysis did achieve high predictive accuracy. This example of supervised machine learning is particularly useful for pattern recognition in highly complex data sets, such as the gut microbiome (40,41). Obviously, as mucosal biopsies are not likely to be routinely harvested in the follow-up of IBD patients, faecal samples have substantial advantages as biomarkers of both disease treatment and activity.

### Healthy Microbial Core

In the present study, microbial signatures of IBD subjects were compared to those of a previously investigated cohort of 61 healthy Dutch children (21). In that study, a shared microbial core, independent of age, and dominated by a limited set of species primarily belonging to the phylum *Bacteroidetes* was identified. Changeability of the core microbiota in children IBD and its tendency to repopulate the faecal microbiome upon achieving remission, underscored that a healthy (non-inflammatory) state may be reflected by an intact core microbiome. This suggests that dysbiosis is primarily characterized by absence of core microbiota, which may be structural in IBD patients, and possibly influenced by genetic or environmental factors. Notably, the observation of an increased *Escherichia coli* abundance at diagnosis of both CD and UC seems to support the keystone-pathogen hypothesis, indicating that certain low-abundance microbial pathogens can induce gut inflammation by manipulation of the commensal normal microbiota into a dysbiotic state (16). Our observation that *Eubacterium spp*. were less abundant in the IBD group compared to the healthy controls is in concordance with findings from previous studies (42,43,44).

Our results are in line with previous observations that de novo paediatric IBD is strongly associated with microbiota alterations. The revised Koch’s postulates need to be fulfilled to ascertain whether this relationship between IBD and bacteria is causal or not. These include that microbial nucleic acids from putative pathogens should be present in most cases of a certain disease and the copy number of these pathogen-associated nucleic acid sequences should decrease or become undetectable upon disease recovery (45). We found that IBD seems to be predominantly associated with a loss of presumably ‘beneficial’ microbes, rather than the introduction of specific pathogens, implicating that IBD, and probably changes within complex microbial niches during chronic inflammatory diseases in general, do not naturally fit these postulates. However, the observed tendency that these healthy core microbes seemed to increase in abundance upon achieving clinical remission of IBD may suggest a causal relationship. It could therefore be questioned whether this phenomenon, a ‘loss of beneficial microbes’ instead of introduction of pathogens, may serve as additional criterion to establish a causal relationship between disease and microbes in chronic inflammatory diseases.

### Microbial Dynamics

We found that treatment-naïve microbiome profiles carry potential biomarkers to discriminate de novo (paediatric) IBD from healthy state. Microbiota has been previously suggested to serve as potential biomarker for follow-up of disease activity, prediction of remission state and therapeutic responses to, for example, anti-tumour necrosis factor (TNF)-α (13,20). After six weeks of treatment, we observed that IBD subjects could still be discriminated from controls, although discrimination between CD and UC was lost, and there was no difference in the overall abundance profile between remission and non-remission. Admittedly, the follow-up period in this study was limited. Another explanation may be that we used PGA scores instead of endoscopic evaluation to evaluate disease activity. The apparent lack of association between microbiota profiles and disease activity in this study hampered their potential as a clinical tool to monitor disease activity, at least during the first episode following initiation of therapy. In addition to compositional differences, we found that over periods of weeks and months the gut microbiota of IBD patients, especially UC subjects, was more unstable over time compared to healthy controls, irrespectively of whether patients were in clinical remission. This finding is in concordance with findings reported in adult IBD patients (46).

#### Exclusive enteral nutrition

Although its mode of action remains largely unknown, a profound impact of EEN on gut microbiota composition and metabolic activity, related to improvement in disease activity, has been described in several studies (35,47). Until now, however, it has not been possible to infer a causative association between EEN induced specific microbial alterations and improvement of disease activity. In the largest study so far exploring associations between gut microbiota and colonic inflammation during EEN, microbiota of 23 CD patients had a broader functional capacity compared to healthy controls, while, unexpectedly, microbial diversity decreased during EEN (18). In our study, we did not observe significant longitudinal differences in diversity indices during EEN course, both in responders as in non-responders. Microbial species abundance at diagnosis had only limited predictive power regarding EEN success at t6. However, in a recent study, comparing the gut microbiome in repeated stool samples of 19 treatment-naïve paediatric IBD subjects and 10 healthy controls, 76.5 % accuracy was obtained for prediction of responder status using pre-treatment microbiome (13).

The strengths of our study are its prospective, multi-centre design with strict standardised sampling and analysis of the microbiota and the relatively large number of patients and a unique control set of samples from healthy children. The inclusion of multiple (longitudinal) samples from controls allowed us to take the substantial physiological variability characterizing healthy state into account in the comparison between microbiota profiles of IBD and controls. The inclusion of treatment-naïve patients allowed us to limit the risk of medication-induced type I error at t0. This study also has some limitations. The follow-up period was relatively short and no endoscopy was routinely performed during the follow-up period to obtain an objective component to assess disease activity. Furthermore, in UC patients and a limited number of CD children who were not prescribed EEN, dietary intake was not assessed during the collection period. However, study and control groups were likely to have comparable diet habits because of a comparable geographic and cultural background. The IBD microbiota profiles were compared to those of an existing cohort of healthy children, which may possibly lead to bias by batch effect. However, samples of both groups were selected in a similar time interval and collection, storage and processing was performed using identical protocols. Mean age of the controls differed from the IBD subjects. In our previous study on microbiota composition, diversity and stability in healthy children, we observed no statistically significant differences between different age groups (2-8 y, 8-13 y, 13-18 y). We therefor believe this has not significantly influenced outcome.

In conclusion, faecal microbiota profiles of children with de novo CD and UC can be discriminated from HC with high accuracy. Therefore, microbial profiling seems to be an effective complementary technique in the diagnosis of paediatric IBD. Both abundance and classification signatures appear to be predominantly driven by a decreased abundance of species which constitute the microbial core in the healthy state, in which *Bacteroidetes* dominate, rather than by the introduction of pathogens. Whether preventive or therapeutic strategies aimed at maintenance or restoration of this core could influence disease activity, needs to be investigated in future studies.

---

**Supplemental Figure 1**. PCoA displaying all IS profiles (all phyla together) samples from Crohn’s disease (A; blue), ulcerative colitis (B; pink) and controls (red) at all time-points. Samples from each individual over time are connected by arrows, from baseline t0□t1□t3□t6 weeks. It can be noted that intra-individual distance of the controls is shorter compared to both ulcerative colitis and Crohn’s disease, indicating that intra-individual stability of controls is higher compared to IBD subjects

**Supplemental Figure 2:** ROC curves. AUC refers to area under the ROC curve. GRR represents the group-regularized logistic ridge. This plot compares performances for the regular logistic ridge regression, GRR, and GRR with post-hoc variable selection. The regular logistic ridge that entertains all retained bacterial features has an AUC of approximately .9. The analogous GRR has a comparable AUC. The question then becomes if this performance can be retained when performing post-hoc variable selection: i.e., can the model be simplified without tarnishing predictive performance? The AUC of the GRR with post-hoc variable selection amount to .87 approximately. Hence, the selected features give a performance comparable to the full bacterial feature-set.

